# Alternating regimes of motion in cell motility models

**DOI:** 10.1101/2019.12.30.891093

**Authors:** Nara Guisoni, Karina I. Mazzitello, Luis Diambra

**Affiliations:** Instituto de Investigaciones Fisicoquímicas Teóricas y Aplicadas (INIFTA), Universidad Nacional de La Plata - CONICET; Instituto de Investigaciones Científicas y Tecnológicas en Electrónica, Universidad Nacional de Mar del Plata - CONICET; Centro Regional de Estudios Genómicos, Universidad Nacional de La Plata - CONICET

## Abstract

Cellular movement is a complex dynamic process, resulting from the interaction of multiple elements at the intra and extra-cellular levels. This epiphenomenon presents a variety of behaviors, which can include normal and anomalous diffusion or collective migration. In some cases cells can get neighborhood information through chemical or mechanical cues. A unified understanding about how such information can influence the dynamics of cell movement is still lacking. In order to improve our comprehension of cell migration we consider a cellular Potts model where cells move actively in the direction of a driving field. The intensity of this driving field is constant, while its orientation can evolves according to two alternative dynamics based on the Ornstein-Uhlenbeck process. In the first case, the next orientation of the driving field depends on the previous direction of the field. In the second case, the direction update considers the mean orientation performed by the cell in previous steps. Thus, the latter update rule mimics the ability of cells to perceive the environment, avoiding obstacles and thus increasing the cellular displacement. Our results indicate that both dynamics introduce temporal and spatial correlations in cell velocity in a friction coefficient and cell density dependent manner. Furthermore, we observe alternating regimes in the mean square displacement, with normal and anomalous diffusion. The crossovers between superdiffusive and diffusive regimes, are strongly affected by both the driving field dynamics and cell-cell interactions. In this sense, when cell polarization update grants information about the previous cellular displacement decreases the duration of the diffusive regime, in particular for high density cultures.

## I. INTRODUCTION

Cell motion plays a key role in many physiological processes including tissue morphogenesis, wound healing, immune and inflammatory response. It is known that the movement of cells is strongly influenced by cell-cell interactions, which grants a wide spectrum of behaviors. Cell motion can be categorized in terms of its external environment, which can present directional asymmetry in response to a chemical stimulus, or be isotropic, without a preferred direction. In this sense, the single cell tracking technique provides substantial evidence that in the absence of chemotactic cues, cells perform a persistent random walk, which has been modeled by the Ornstein-Uhlenbeck (OU) process [1]. In the case of directional asymmetry of the environment the cell is said to perform taxis and cell movement has been widely modeled using the Keller-Segel diffusion equation [2]. In addition to this categorization, cell motion can refer to the movement of individual cells with or without neighbors [3, 4], or to a cell population acting as an aggregate [5–7].

Experimental results focused on individual movements share characteristics of Brownian particles [8, 9]: exponential decay of the velocity autocorrelation function (ACF) and linear growth with time of the mean square displacement (MSD), at large time scales [10]. These features can be explained by the OU process, or equivalently, the conventional Klein-Kramers description [1, 11]. However, there exist cell motions without chemotaxis that do not follow the OU process, for example, movements of epithelial cells and aggregates of Hydra cells, reported as anomalous diffusion processes [5]. Also human fibroblasts and keratinocytes move in a manner that contradict the OU process [5]. Similarly, Takagi et al. have reported different cell movement behaviors with anomalous diffusion for Dictyostelium cells in different physiological conditions [12]. These experiments fit well with a generalized Langevin model that includes a memory kernel for cell velocity [13]. Furthermore, long-term analysis of MDCK cells has revealed a superdiffusive behavior, in absence of external cues [4, 14]. These findings show that cell movement contains a more complex dynamics than the persistent random walk, which can be explained by the fractional Klein-Kramers equation [4]. The latter can be considered as a phenomenological approach, able to describe anomalous diffusion in terms of very general physical mechanisms. However, it has limitations to indicate which biological ingredients lead to an anomalous behavior. Thus, alternative modeling that allows to get biological insight by testing different hypotheses becomes really interesting.

Recently, we have introduced a reorientation model based on the cellular Potts framework [15]. In this model, cell movement due to a driving field, which direction changes following a discrete version of the OU process, was considered. In contrast, previous models have applied the OU process on the velocity vector, leading to white-noise fluctuations on the direction angle [8, 9, 16, 17]. It is known that when the orientation angle fluctuates without correlations (i.e., in the absence of the friction term) the system exhibits Brownian motion [16, 18]. However, when the friction term is present, we found that high density cultures exhibited a double-exponential for the velocity ACF, in contrast to an exponential characterizing the Brownian motion of low density cultures. For both densities the MSD behaves as a persistent random walk model for the time scale studied [15]. These results suggest that the complex behavior of cell motion can be consequence of the intrinsic feature of the movement, but also of cell-cell interactions.

In this paper, we are particularly interested in understanding the interplay between the neighborhood information gathered by the cell in previous displacement and cell motility over large time scales. To address this question, the cellular Potts model (CPM) introduced in [15] with two types of dynamics for the orientation angle of cell displacements are compared. Firstly, we consider a “naive” implementation where the new orientation is related to the field direction operating in the previous step, regardless the cellular direction of displacement. In the second case, the update direction depends on the mean orientation performed by the cell in the previous steps. This implementation of the field direction update grants a sort of feedback mechanism, because at each time step the angle of the driving field is influenced by the recent cell history, taking into account interactions between cells blocking and deviating from their original orientations. We compare both actualization models analyzing the MSD, the temporal and spatial correlations of cell velocity and the average distance traveled by a cell during the time interval that the driving field is operating, at low and high density cultures.

## II. THE MODEL

The CPM is a modified Potts model which includes different terms of energy that become it able to reproduce some biophysical properties of cells, such as deformations of cell membrane, adhesion and motility in an excluded volume manner. In the model, at each site of the lattice a spin *σ*_*i*_ = 1, …, *Q* is assigned, and cells are represented by domains with the same spin, thereby if *σ*_*i*_ = *M*, with 1 ≤ *M* ≤ *Q*, it belongs to the cell labeled as *M*. The dynamics of the model are governed by the Hamiltonian, or energy function, which guides the cell behavior by distinguishing the low energy configurations (or favorable) from the high energy ones. The Hamiltonian is constituted by a term corresponding to the sum of all surface energies, responsible for cell-cell adhesion properties. However, to keep the cells without that they be broken or disappear, additional terms in the Hamiltonian are needed. Thus, the energy function considered here is given by

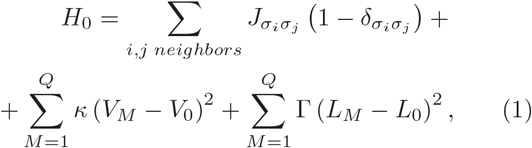

where 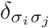 is the Kronecker delta and the first sum is over all neighboring site pairs, representing the boundary energy of the interacting cells. The second and third terms in Eq. (1) correspond to the energy costs for cells to deviate from the preferred volume *V*_0_ and perimeter *L*_0_, respectively. The presence of a medium, which interacts with the cells, is also considered. In this way, the medium has spin variable *σ*_*i*_ = 0, with no target area or perimeter. The adhesion constant between different cells is denoted by *J*_*cell-cell*_ whereas between cells and medium is *J*_*cell-medium*_. Further, to consider cell motility preferentially along the direction of a driving field 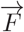 an additional term should be added to the Hamiltonian Eq. (1) [15, 19], as we will see below.

The system evolves using Monte Carlo dynamics. In order to obtain a new configuration a lattice site is randomly chosen and if it belongs to the boundary of the cell, this site copies the spin value of one of its neighboring cells as a trial. The variation of energy in a proposed trial configuration is given by

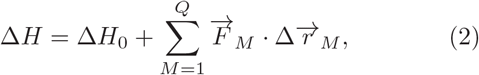

where Δ*H*_0_ is the change of energy related to Eq. (1), 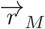 denotes the displacement of the center of cell *M* and 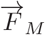 is the driving field acting on cell *M*. The acceptance, or not, of a new configuration is given by the Metropolis prescription: the trial is accepted with probability 1 if it decreases the value of energy, Δ*H* ≤ 0, or with the Boltzmann factor 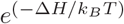 if it increases the energy (Δ*H* > 0), where *k*_*B*_*T* parameterizes the intrinsic membrane motility. The unit of time, a Monte Carlo step (MCS), is defined as *N* trials of movement, being *N* the number of spins in the lattice.

The driving field is characterized by a direction, denoted by Θ, and an intensity *F*. We consider that the intensity *F* is constant over time and the same for all cells. However, the direction of the driving field operating over cell *M*, Θ_*M*_, is actualized according to OU process, *d*Θ_*M*_(*t*) = −λΘ_*M*_ (*t*)*dt* + *σdW* (*t*), where λ is the friction coefficient (0 ≤ λ < 1), *σ* determines the magnitude of the fluctuations and *dW* (*t*) denotes the Wiener process. For our Monte Carlo simulations, it is necessary to use a discrete version of this stochastic differential equation, which can be identified with a first-order autoregressive process, as follow:

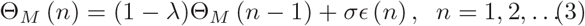

where *ϵ* (*n*) is a white noise with zero mean and unit variance 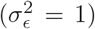 and *n* is the discrete time. λ and *σ* were defined previously.

Note that it is not mandatory that the cell displacement has the same direction of the associated driving field, due to both cell-cell interactions and stochastic fluctuations. Thus, the angle of the driving field Θ_*M*_ is not necessarily equal to the direction of the cell movement, which will be denoted by *α*_*M*_. If the previous direction of the cell displacement is considered in the actualization of the driving angle we have a positive feedback, which mimics the situation in which the cell produces its own chemotactic signal. This aspect was taken into account previously by other authors in Potts like models [6, 20, 21]. In that way, Kabla considers that the motile force is oriented along the mean velocity of the cell over its past time steps, without friction nor noise [21]. For Szabó et al. [6] the change in cell polarization is proportional to a spontaneous decay respect to its previous value and a reinforcement from cell displacement direction during the time step considered. In order to take into account this feedback loop, an alternative way to update the angle of the driving field Θ_*M*_ is:

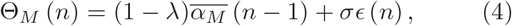

where 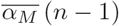 is the mean displacement angle over the last *τ* MCS, and the other parameters are the same as in Eq. (3). Thus, according to Eq. (4) the angle of the driving force depends on the earlier displacements of the cell. Differently from previous formulations [6, 20, 21], our proposal for the feedback loop takes into account fluctuations. Besides, an advantage of Eq. (4) is the possibility of a direct comparison with Eq. (3): the only difference between them is the dependence on the mean cell polarity 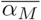 instead of on the previous direction of the driving field *θ*_*M*_.

For both actualization procedures, the initial direction of the cell *M*, Θ_*M*_ (0), is chosen randomly between [0, 2*π*]. Θ_*M*_ evolves independently of the field operating in other cells. The updating time in Eqs. 3 and 4, *n*, is different from the time of actualization of cell configurations. In particular, at each time step the direction of the driving field for each cell *M* changes with probability 1*/τ* according to Eqs. 3 and 4. Thus, the change in the directions Θ_*M*_ and *α*_*M*_ occurs at a mean time *τ* independently of the direction of other cells.

## III. RESULTS

For all simulations in this paper, we used the following fixed parameter values *J*_*cell-cell*_ = 0.1, *J*_*cell-medium*_ = 0.01, Γ = 0.2, *κ* = 1, *σ* = *π/*3, *F* = 10, *T* = 2 and *τ* = 10. The density *ρ* is defined as the ratio between the area occupied by the cells and the medium. In this way, we calculate the number of spins with *σ*_*i*_ ≠ 0 related to the total number of spins, since the medium is identified by *σ*_*i*_ = 0. Low and high density simulations correspond to *ρ* = 0.2 and *ρ* = 0.9, respectively. Also, we considered along the paper three different values of friction coefficient λ = 0.01, 0.05 and 0.10. We used periodic boundary conditions and a square lattice of size 1024 × 1024 sites. More details about initial conditions and thermalization can be found in [15].

In order to characterize the movement of a cell population we calculate the mean-squared displacement (MSD) as 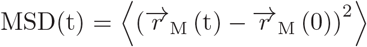, where the average is taken over all cells of the simulation between a common starting point at *t* = 0 and the actual positions at time *t*. According to Fig. 1, the MSD presents two or three regimes in the time scale considered, depending on the value of λ and on the updating rule used. For λ = 0.01 and the OU actualization (Eq. 3), we can see that at short times the MSD is almost ballistic and after that it resembles a random walk, regardless the density. For the other situations, the MSD has three regimes: it is almost ballistic at short times, diffusive at intermediate time scale and superdiffusive at long times. When the direction of the driving field is actualized by using OU with feedback (Eq. 4), the crossover between the random walk behavior and the superdiffusive regime at long times occurs previously for λ = 0.10 than for 0.01. On the other hand, when the OU actualization is used, this crossover is present for λ = 0.10 but not for 0.01. Consequently, the duration of the diffusive period is shorter when the friction coefficient is higher (same actualization model and different values of λ) and when the feedback is present (same value of λ and different actualization models). In order to understand these results, let us discuss the meaning of the different regimes observed for the MSD along the different time scales. For all cases, the almost ballistic short-time behavior is related to the persistence time of the driving force, *τ*. In fact, in a previous paper [15] we shown that the temporal behavior of the MSD scales with *τ*, for the OU actualization. The diffusive behavior of the MSD, also present for all cases shown in Fig. 1, is the result of the fluctuations in the direction of the driving field, since Eqs. 3 and 4 are stochastic equations. However, the actualization of Θ is not completely random since the friction coefficient λ introduces correlations in the successive directions of the cells displacements, whose influence is observed at long times. This effect is particularly evident when using Eq. 4, since the presence of feedback rises the correlations, and therefore favors the anomalous diffusive behavior.

**FIG. 1.**
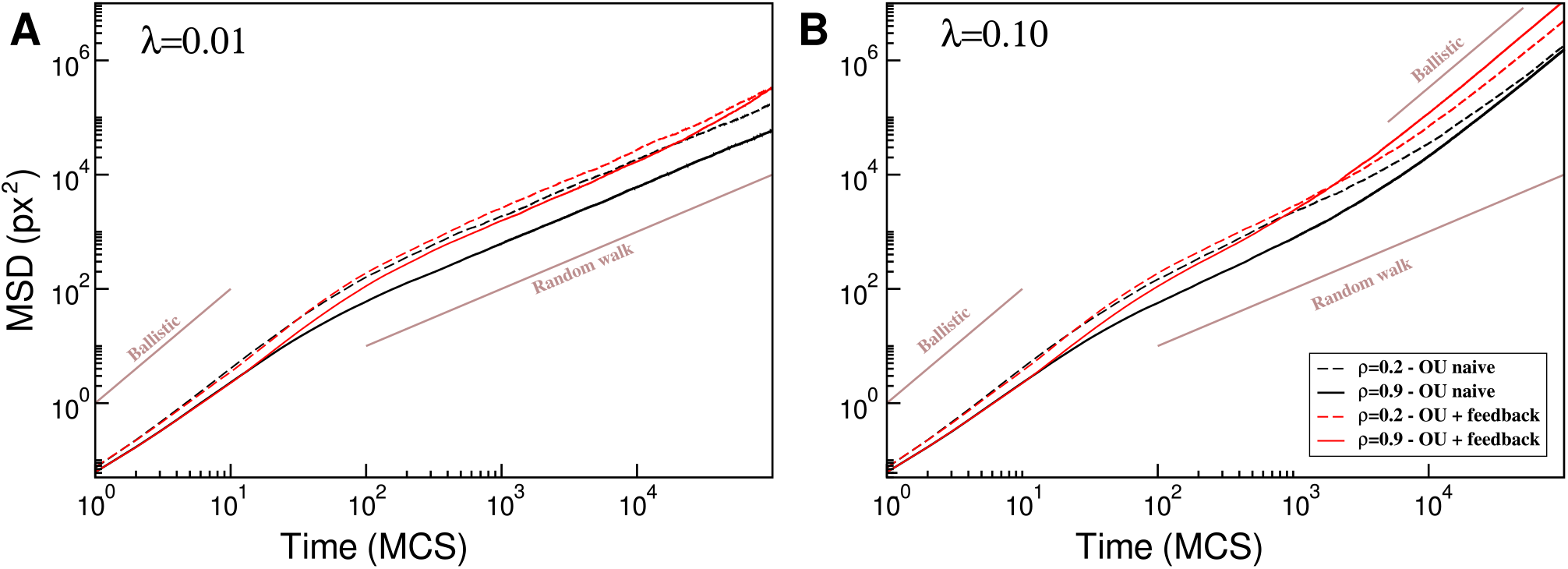
Log-log plot of the mean-squared displacement, msd, *vs.* time, *t*. Direction of the driving field actualized according to Ornstein-Uhlenbeck (OU naive) and Ornstein-Uhlenbeck with feedback (OU + feedback), λ = 0.01 (A) and λ = 0.10 (B), and ρ = 0.2, 0.9 (dashed and continuous line, respectively). Lines with slope equal 2 (Ballistic) and 1 (Random walk) are shown for the sake of comparison.

Also, from Fig. 1, the MSD obtained from the angle updating rule with feedback is greater or equal than that found with the OU actualization. This result suggests that feedback helps cells avoid collisions with another cells, making the movement more effective. Finally, we discuss the effect of density on the MSD, starting with the model of OU actualization. Low density configurations have higher or equal MSD than the obtained for high density, for both values of λ (see black lines in Fig. 1). In fact, a lower MSD for high density cultures is expected, since crowded cell cultures usually disturb the movement of cells. The same behavior related with density can be observed at short and intermediate times when the feed-back updating rule is used. However, at long times, the high density culture presents higher MSD than the low density one, for λ = 0.10 and feedback update. This inversion in the MSD suggests that the feedback gives rise to a spatially coordinated movement of the cells in high density simulations.

The temporal behavior of MSD scales with time as MSD(t) ∼ t^*β*(t)^, where the exponent *β* characterizes the different regimes observed. In that way, a ballistic behaviour is associated with *β* = 2, whereas normal diffusion presents *β* = 1. The logarithmic derivative of the MSD allows the calculation of *β* as 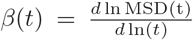. Similar measurements were used to study cell migration and intracellular transport [22] from both experiments and models, and for a simple model that mimics the diffusion of a particle in an anisotropic amorphous material [23]. Fig. 2 shows the behavior of *β* as obtained from the two angle updating rules, for low (dashed lines) and high (solid lines) densities and different values of λ. *β* was computed using a time-sliding window, the size of which depends on *t*. At the short-time scale the exponent *β* corresponds to anomalous diffusion. At this scale, we can note that for the OU actualization, low density cultures present higher *β* than high density. This aspect is less evident when the feedback mechanism is considered, since *β* for low and high density cultures presents almost the same value in the range [30, 100] MCS. Besides, at short-time scale it can be seen a slight increase in *β* for the update with feedback. These results indicate that the feedback makes cell movement more efficient, particularly for high density cultures, as discussed before. At intermediate time scale, *β* decreases and the MSD tends to exhibit a diffusive behavior. Particularly, for the OU actualization and low λ-values, the diffusive behavior is observed at intermediate and long-time scales. For the other conditions of Fig. 2, the transition between short-time and long-time superdiffusive regimes is so tight that the exponent *β* = 1 is almost not reached, but will be referred to as a diffusive regime. The duration of this diffusive regime decreases with λ (for the same angle updating rule) and with the feedback (for the same value of λ), as pointed out with Fig. 1. Also, for the OU actualization and low λ-values, *β* is independent of the density at intermediate and long-time scale. For the other conditions in the same time scales, the high density cultures present higher *β* than low density ones. Besides, from the behavior of *β* at long-time scale when the feedback is considered (Fig. 2-B), we can suppose that for a larger time scale (that is for *t* ≥ 10^5^ MCS) *β* will continue to grow until it reaches *β* = 2. In fact, we can expect the same behavior for Fig. 2-A, since for both actualization rules there is correlation in the Θ update. As we have discussed previously, the correlation comes from the friction term being reinforced by the feedback. Cell-cell interactions also increase correlations at intermediate and long-time scales.

**FIG. 2.**
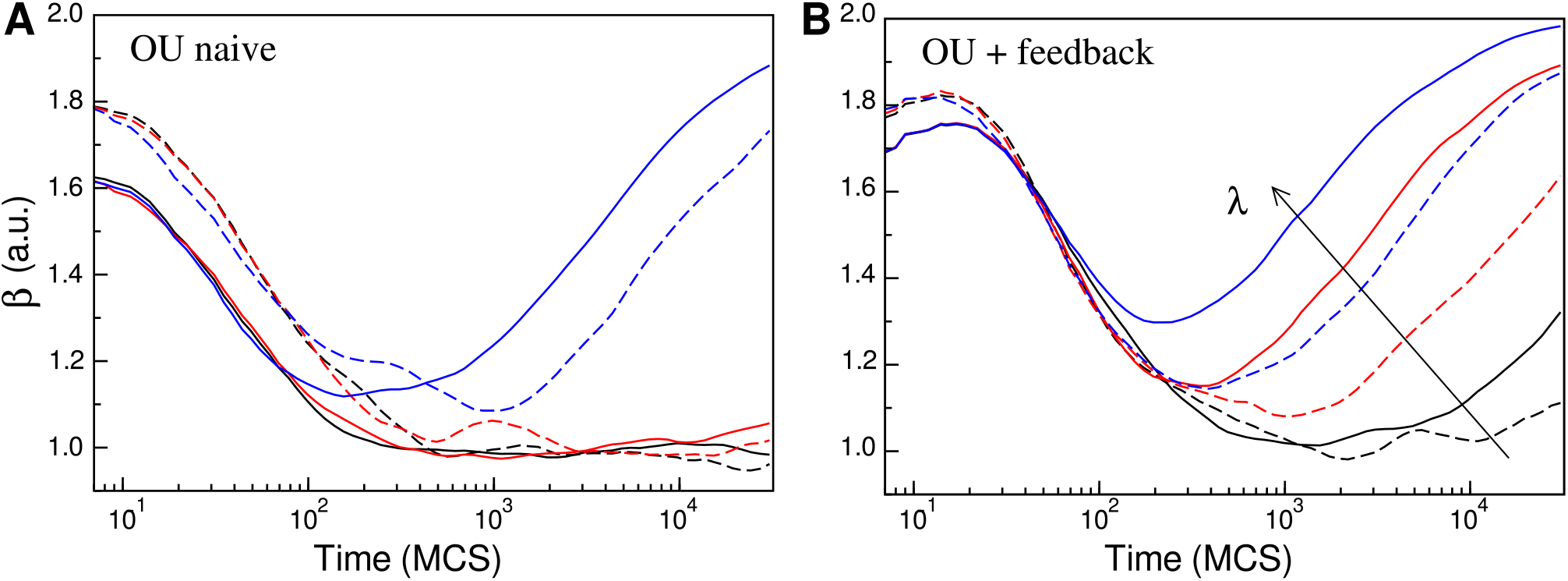
Logarithmic derivative of the MSD, *β*(*t*), as defined in the text, calculated from data shown in Fig. 1 and additional data. Direction of the driving field actualized according to OU naive (A) and OU with feedback (B) and different values of λ = 0.01, 0.05, 0.10 (dark, red and blue, respectively) and *ρ* = 0.2, 0.9 (dashed and continuous line, respectively). We used a time-sliding window, the size of which depends on time. The interval between successive measurements is equal to [1.775^*k*^, 1.775^(*k*+4)^] and *k* = 1, 1.2, 1.4,…, 16, since the MSD data was considered until *t* = 10^5^ MCS.

The velocity ACF is another important tool to characterize cell movement. It is defined as *C*(*t*) = *Z*(*t*)*/Z*(0) where 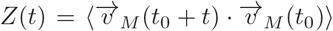 [24], and 〈…〉 indicates the average over all cells and over *t*_0_. The cell velocities are defined as 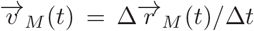, where 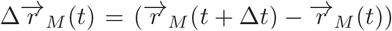, and Δ*t* = 1 MCS. Fig. 3 shows that the velocity ACF obtained for the angle updating rule with feedback (Eq. 4) is always higher than the one obtained for the naive angle update (Eq. 3), regardless the values of λ or *ρ*. In fact, if cells are more successful in avoiding collisions with another cells we expect a higher ACF. In Fig. 3, for the OU actualization, high density cultures have smaller ACF than low den sity cultures, for both λ = 0.01 and 0.05. This result can be understood by the fact that a crowded neighborhood usually disrupts the movement of the cell. However, when the feedback update is considered, this relation is inverted: high density cultures have a higher ACF than low density cultures, independent of the value of λ. Therefore, we can conclude that the feedback makes cell movement more efficient mostly for high density cultures. Actually, when the update with feedback is used in high density cultures there is a competence between two effects: on the one hand a crowded environment hinders cell movement, and on the other hand, the feedback promotes it. But the OU update has only the first effect. Because of that, the difference in the velocity ACF between the two update rules is more evident for high density cultures.

**FIG. 3.**
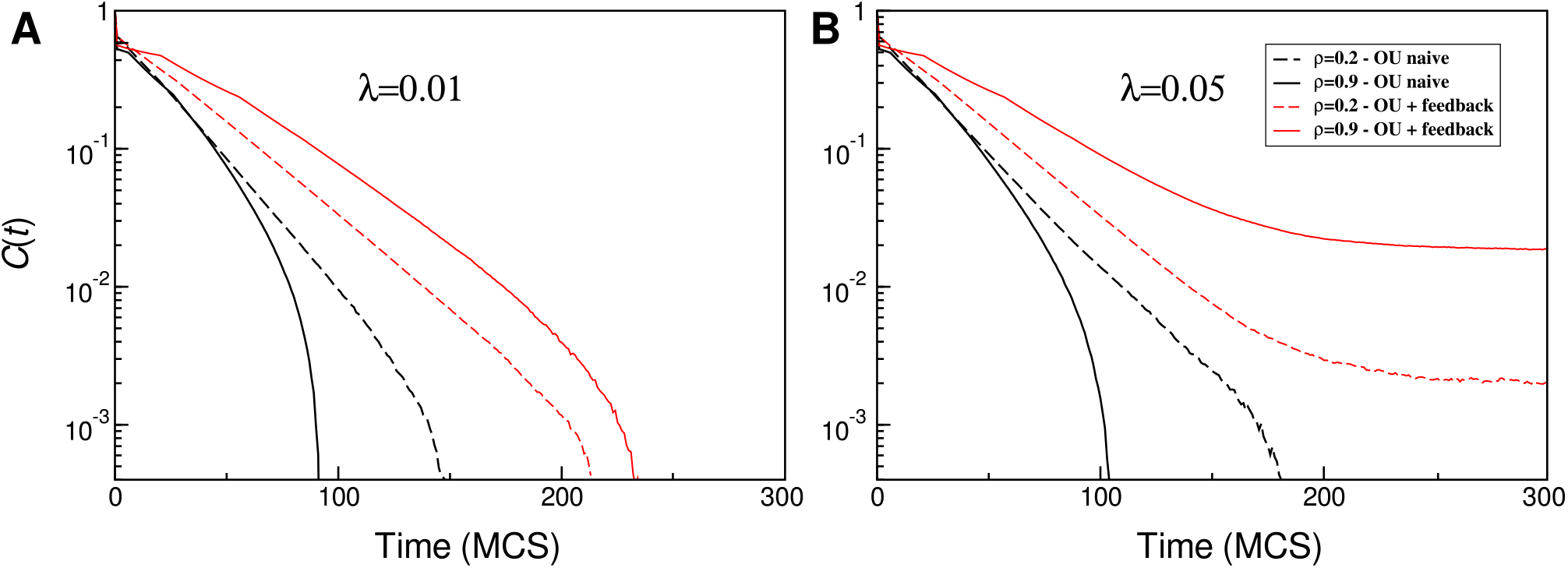
Semi-log plot of the velocity ACF function, C(t), *vs.* time, *t*. Direction of the driving field actualized according to OU naive and OU with feedback, for different densities, *ρ* = 0.2, 0.9 (dashed and continuous line, respectively), and friction coefficients, λ = 0.01, (A) and λ = 0.05 (B). C(t) was averaged over all cells in the simulations for 6×10^4^ MCS and over 1 (*ρ* = 0.9) and 3 samples (*ρ* = 0.2).

Also, Fig. 3 shows that the ACF is greater for λ = 0.05 than for λ = 0.01, regardless the density or the update model. This result indicates that the friction coefficient enhances the correlation in cell movement, as discussed before. Finally, for the OU with feedback update and λ = 0.05 the ACF presents a very slow decrease at long times, consistent with an algebraic decay, independent of the density. For the other parameters considered in Fig. 3 the ACF goes to zero before *t* = 250 MCS.

The spatial correlations of the cell velocities are also used to study cell movement. It is defined as 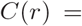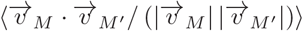, with 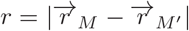 the distance between mass center of cells *M* and *M* ^′^. When comparing the two angle-update rules, we can see from Fig. 4 that *C*(*r*) is always higher when the update rule with feedback is used, as also observed for the ACF (Fig. 3). As discussed in a previous work [15], the peak close to the typical diameter of the cell (∼ 16 pixels) is related to anti-correlated velocities of cells that travel in opposite directions. At intermediate and long distances (*r* ≳ cell size) cells in high density configurations are more correlated, for both actualization rules and λ−values. Besides, *C*(*r*) in high density simulations approaches zero very slowly, as *r* increases, in particular for λ = 0.10 and when using the feedback update rule (Fig. 4-B). These results indicate that cell-cell contact induces long-range spatial-correlation of cell velocity and that this effect is enhanced by both the feedback and the friction coefficient. Also, the difference in *C*(*r*) for low and high density cultures is much higher in the case of update with feedback. Therefore, the effect of the feedback raising spatial correlations is greater for the crowded cultures, as observed for the temporal correlations in Fig. 3. These results could indicate a coordinated movement of cells in high density cultures as a consequence of the feed-back mechanism.

**FIG. 4.**
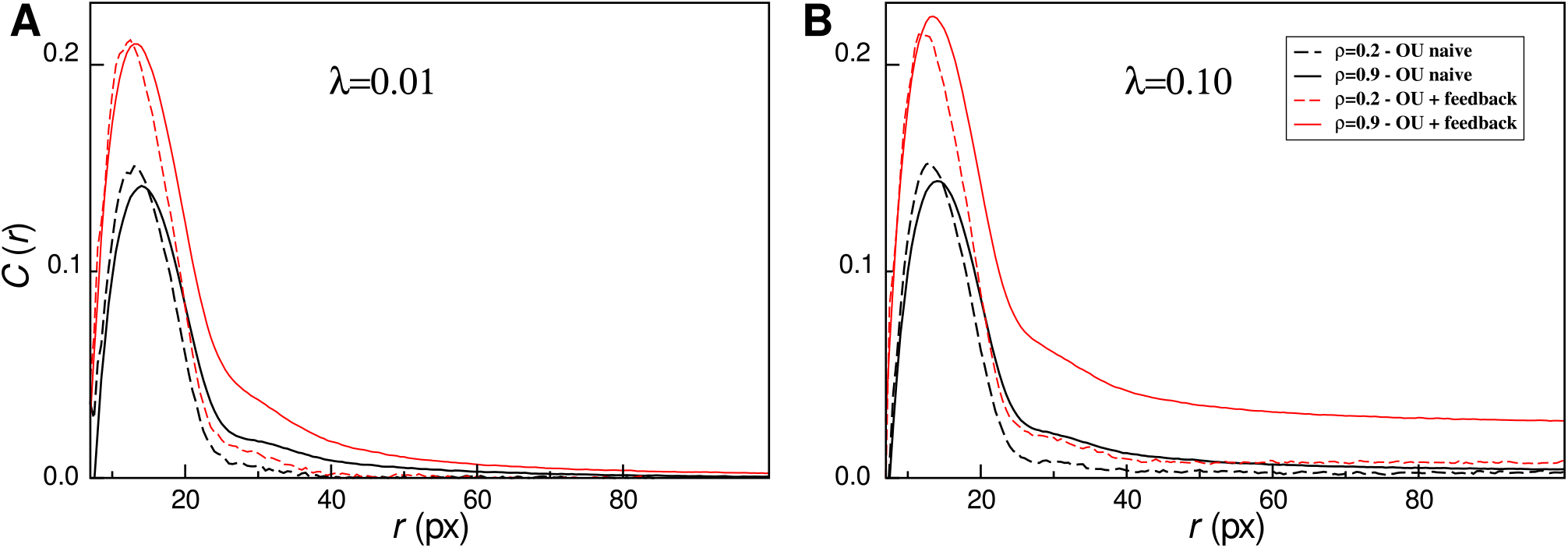
Spatial correlation function of the velocities, *C*(*r*), as a function of the distance *r* between cell pairs. Direction of the driving field actualized according to OU naive and OU with feedback, for different densities, *ρ* = 0.2, 0.9 (dashed and continuous line, respectively), and friction coefficients, λ = 0.01, (A) and λ = 0.10 (B). Data were obtained by averaging over 10^5^ MCS.

In order to get insight about the effect of the feedback introduced by the rule Eq. 4 on the cell movement, we define the average distance *D̅* traveled by a cell during the time interval that the driving field is operating in a given direction. Mathematically, it is defined as

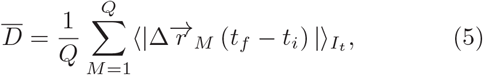

where 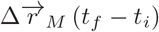 is the cell displacement from the time the driving field starts operating, *t*_*i*_, until the time of the next direction change, *t*_*f*_ (with *t*_*f*_ − *t*_*i*_ ∼ *τ*), 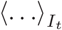 indicates the average over all time intervals *I*_*t*_ = [*t*_*i*_, *t*_*f*_], and the sum runs over all cells on the substrate. This magnitude gives information about the cell movement only in the short-time scale (∼ *τ*, i.e., during the first almost ballistic regime), for that reason it is nearly independent on λ-values, as we can see in Fig. 5. Further, it is expected that cells can move much more in low density cultures than in a crowded media, which is also evident in Fig. 4, where *D̅* is much higher for *ρ* = 0.2 than for *ρ* = 0.9. Fig. 5 also establishes a comparison of the magnitude *D̅* for the two angles-updating rules specified by Eqs. 3 and 4. In all cases, the average distance *D̅* reached by the cells when the OU with feedback is acting is greater than when the OU update is applied. In particular, we observe that in the case of a crowded media there is a remarkable increment (of about 20%) of *D̅* when the update with feedback is applied respect to the OU actualization. For the low-density case the increase is only about 3%. These results suggest that the OU with feedback update promotes or increases cell displacement and that this effect is more noticeable in crowded environments.

**FIG. 5.**
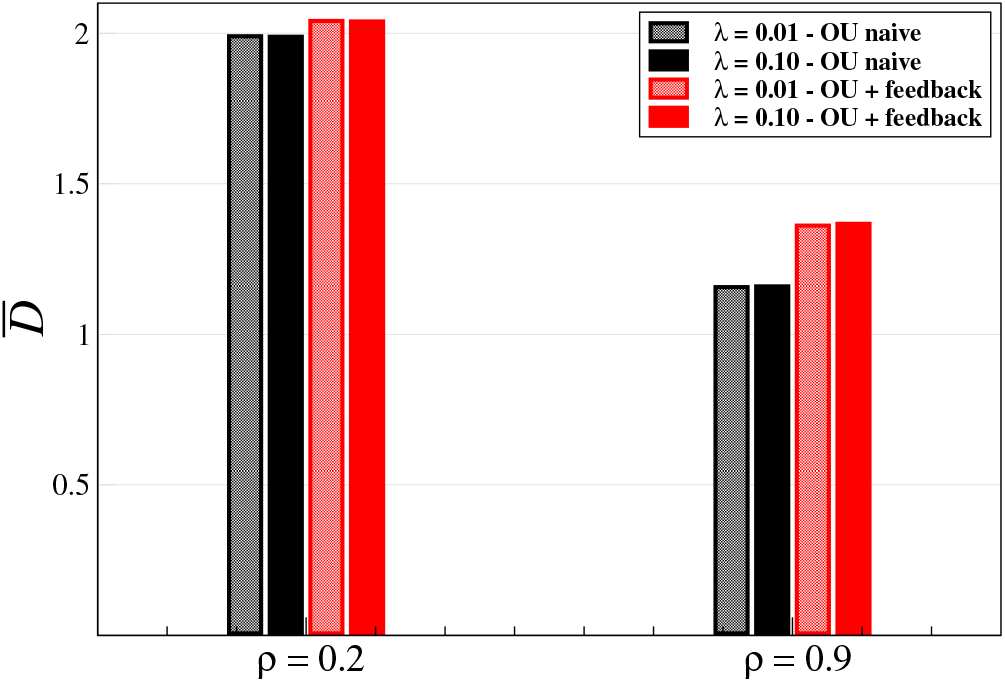
Average distance travelled by a cell during the time interval that the driving field is operating, *D̅*, for densities ρ = 0.2 and ρ = 0.9. Direction of the driving field actualized according to OU naive and OU with feedback (black and red, respectively) and friction coefficient λ = 0.01, and λ = 0.10. *D̅* was averaged over all cells in the simulations until 10^5^ MCS (the first 500 MCS were disregarded). Error bars are smaller than the thickness of the line.

## IV. DISCUSSIONS AND CONCLUSION

Much of the work about cell motility is based on the study of the time behavior of the second moment, the MSD. Thus, a system is considered to exhibit Brownian motion when the MSD increases linearly in time, otherwise it is considered to present anomalous diffusion. However, we usually want to know more about the cell trajectories than simply the second moment [3, 4]. Other features of interest are the distribution and correlations of cell velocity. So, if normal diffusion occurs, the velocity correlation decreases to zero exponentially, or more quickly, while anomalous diffusion could be associated with an algebraic decay of the velocity ACF. The MSD and the correlations can be theoretically derived only from simplified models [13, 16, 17, 22, 25]. This modeling feature is particularly interesting to get insight about the asymptotic behavior of the MSD. On the other hand, for models which include biological ingredients like cell volume and cell-cell interactions, as the CPM-based ones, it is difficult to get analytical expressions. However, these models can add valuable information about cell motion [6, 15, 21, 26]. For example, they are able to test feedback mechanisms from the neighbourhood at different spatial scale in an excluded volume schema [21, 26]. In addition, Kabla studies different cell dynamics for the collective movement resulting from the balance between adhesion and cell motile forces [21]. Here we study cell movement at long-time scales addressing the effect of cell-cell interactions and neighborhood information gathered by the cell within the framework of a CPM-based model. We consider that cells move actively according to a driving field that has a constant intensity and whose orientation is governed by two alternative OU updating rules. The proposed dynamics for the driving field provide both a persistent random walk feature, which include a sort of angle memory, as well as, a feedback mechanism able to mimics cell behavior with environment perception at local range.

We observed alternating superdiffusive-Brownian regimes for the MSD in the temporal scale considered. The almost ballistic behavior at short time is related with the persistence time *τ* of the driving field [15]. At intermediate time intervals the MSD becomes diffusive due to the random actualization of cell direction. A crossover from a quadratic to a linear regime in the MSD has been previously reported in a model of self-propelled particles, when the diffusive behavior arising from particle reorientation dominates the persistence process [17]. Besides, when the OU dynamics is applied to update particle velocities vector (instead to update the angle direction as was done here) the same crossover is observed in the MSD and the resulting asymptotic regime is Brownian [1, 16]. In addition to this initial crossing of regimes, we found a second crossover between the diffusive behavior and a ballistic regime at large time scales. The second crossover occurs previously for large λ-values, since the stronger the friction term in the OU process, the higher the correlations. Furthermore, we show that this crossover also depends on the angle-update rule used, being favored when Eq. 4 is operating. This result is in agreement with the fact that temporal and spatial correlations are higher when the update rule with feedback is used. Also, our results suggest that the feedback update rule helps cells avoid collisions with another cells, making the movement more effective, and therefore contributing to sup-perdifusion. Finally, the transition to the long-term ballistic regime depends on cell-cell interactions and occurs previously in crowded cultures. Our findings can be related to previous results from a directed random walk model [27]. In this paper, the time of appearance of the asymptotic ballistic regime depends on an anisotropy parameter, which fixes the correlations in the displacement direction: at higher values of this parameter (or stronger anisotropy), there is more correlation and the ballistic regime appears early [27]. In the present work, we show that the appearance of the long time ballistic regime is favored by the friction term λ, by cell-cell interactions and also by the feedback updating rule. All these factors introduce correlations in cell movement, leading to a ballistic motion at long time scales. We also expect the crossover time between the diffusive and the ballistic regime be affected by *σ*, since greater values of *σ* are related to a more stochastic cell movement and should correspond to a longer diffusive period.

In conclusion, the two angle-updating rules used in this paper allow disentangling the effect of two sources of anomalies. Firstly, we observe that cellular motion governed by angle-updating rules with a no-null friction term, presents anomalous diffusion for long times, more specifically a crossover from Brownian to ballistic regimes. On the other hand, the temporal scale of the diffusive regime is shortened when the direction update rule includes a feedback mechanism. The characteristic scale where the crossover between these regimes is evident depends on the velocity correlations, which itself depends on the friction term, the feedback and cell-cell and interactions.

## ACKNOWLEDGMENTS

We acknowledge financial support from the Argentinian Science Agencies CONICET and ANPCyT, and from the UNLP (Universidad Nacional de La Plata).

